# Appelmans Protocol for *in vitro Klebsiella pneumoniae* phage host range expansion leads to induction of a novel temperate linear plasmid prophage *vB_KpnS-KpLi5*

**DOI:** 10.1101/2023.08.05.552120

**Authors:** Nadine Jakob, Jens A Hammerl, Brett E Swierczewski, Silvia Würstle, Joachim J Bugert

## Abstract

Adjuvant therapy with bacteriophage (phage) cocktails in combination with antibiotics is a therapeutic approach currently considered for treatment of infections with encapsulated, biofilm forming, and multidrug-resistant *Klebsiella pneumoniae* (Kp). *Klebsiella* phage are highly selective in targeting a bacterial capsule type. Considering the numerous Kp capsule types and other Kp host restriction factors, phage treatment could be facilitated when generating phages with a broad host range A modified ‘Appelmans protocol’ was used to create phages with an extended host range via *in vitro* forced DNA recombination. Three T7-like Kp phages with highly colinear genomes were subjected to successive propagation on their susceptible host strains representing the capsule types K64, K27, and K23, and five Kp isolates of the same capsule types initially unsusceptible for phage lysis. After 30 propagation cycles, five phages were isolated via plaque assay. Four output phages represented the original input phages, while the fifth lysed a previously non-permissible Kp isolate, which was not lysed by any of the input phages. Surprisingly, sequence analysis revealed a novel N15/phiKO2-like phage genome (*vB_KpnS_KpLi5*) lacking substantial homologies to any of the used T7-like phages. This temperate phage was only induced in the presence of all input phages (cocktail), but not by any of them individually. Induction of temperate phages may be a stress response caused by using multiple phages simultaneously. Successive use of different phages for therapeutic purposes may be preferable over simultaneous application in cocktail formulations to avoid undesired induction of temperate phages. (243)

## Introduction - Results - Discussion

Multidrug-resistant (MDR) *Klebsiella pneumoniae* (Kp) are one of the consequences of inappropriate use of antibiotics in patients and animals and/or their discharge into the environment (One Health approach). As typical ESKAPE(E) bacteria, Kp efficiently arm-up their genomes acquiring genetic material from other bacteria to adapt to environmental and artificial selection pressures (i.e., antimicrobials, biocides). Nowadays, MDR Kp can be found in many places in the environment, including surface water and sewage treatment plants [1,2]. Treatment of human Kp infection is challenging as isolates exhibit various defense strategies, e.g., a large variety of capsule types (>80). In addition, Kp form biofilm matrices, enabling sustained survival in a broad range of environmental and host conditions. In view of high case fatality rates in intensive care patients [3], alternative therapeutic methods must be thoroughly explored [4].

The above-mentioned diversity of Kp capsule types and further restriction factors of Kp lead to the need for a large amount of the very host specific Kp phages to cover all types required for therapy. However, even a phage bank with more than 350 phages with different capsule types is not sufficient to efficiently lyse the respective Kp target population of patients[5].

Thus, a modified ‘Appelmans protocol’ [6,7; Figure 1A], was used to create phages overcoming non-capsular - restriction factors via forced DNA recombination between colinear phages in initially non-susceptible/permissible hosts. Three T7-like phages (*Caudovirales, Autographiviridae, Studierviridae*, Prondovirus spp.) with moderate to high genome colinearity were selected to allow homologous recombination between the phages (Figure 1B): Tun1(Genbank # HG994092.1) [8], EkP2 (WRAIR; Genbank # OQ921102), Muc100 (IMB collection; Genbank # OQ921101). The hypothesis to be tested suggested that genome rearrangements forced by selection on non-susceptible test hosts would overcome additional host restrictions leading to phages able to lyse the test hosts. Eight strains were used, comprising three original hosts strains (Kp 7984 (capsule type K64) from the Military Hospital Tunis, Tunisia, Kp KRI 9 (K27) from a Munich hospital, Germany, and Kp KpA2 (K23) from Walter Reed Army Institute of Research (WRAIR), MD, USA, as well as five Kp isolates with the same capsule types (test hosts), but not susceptible to lysis with the T7 phages, indicating additional restriction factors (Table 1).

**Table 1.**
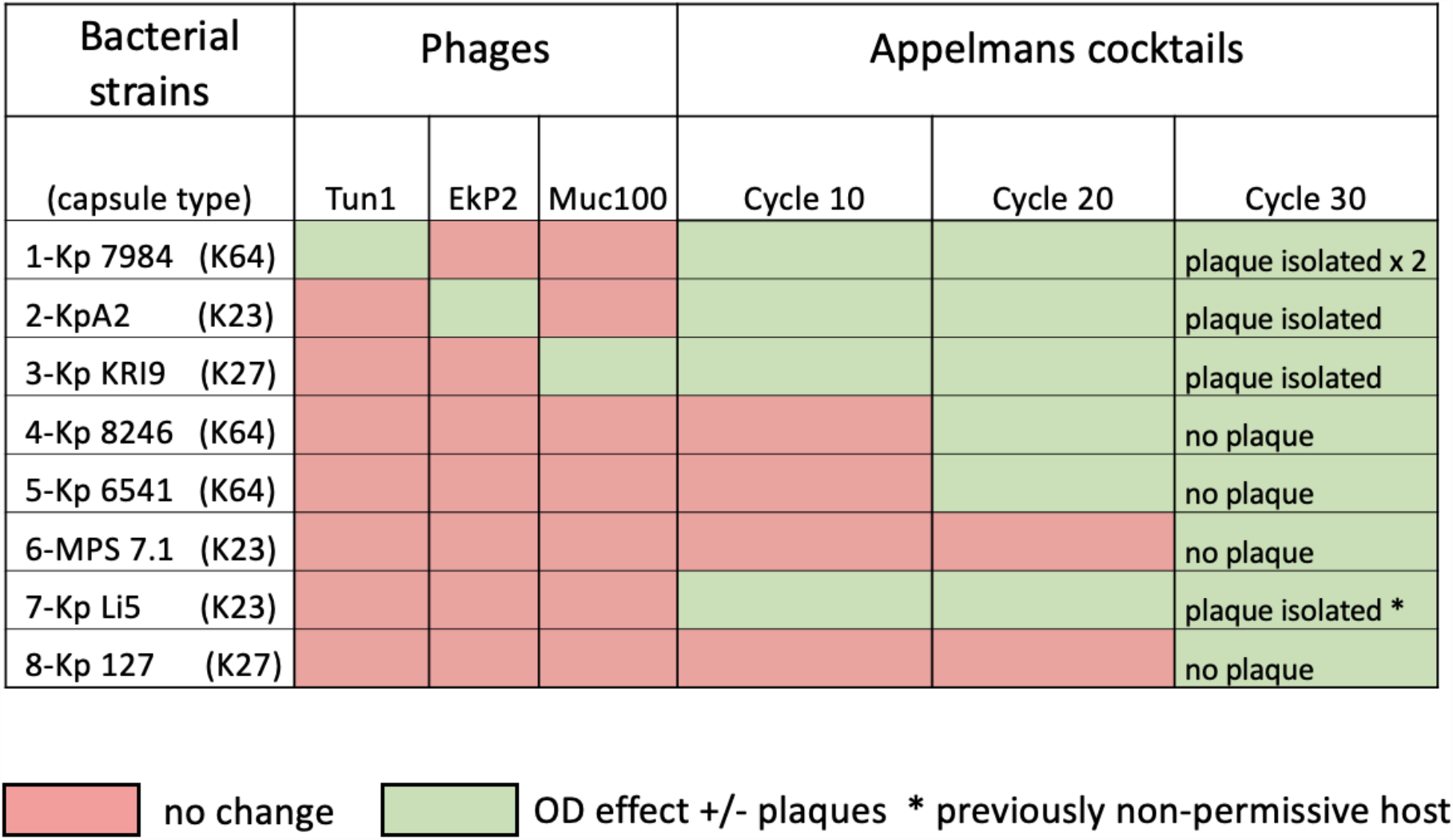
(b/w) Bacterial strains and phages.

**Figure 1.**
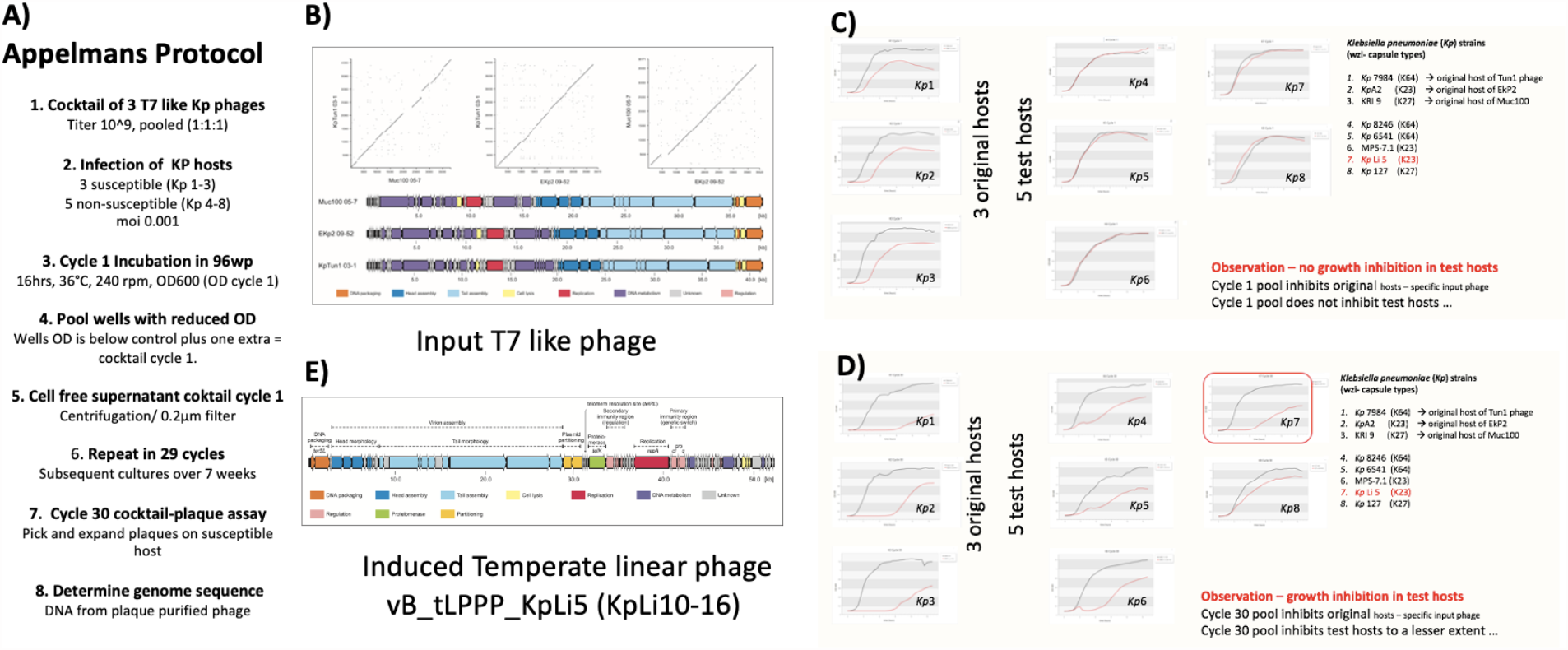
Experimental protocol and results. A) modified Appelmans protocol; B) input phage KpTun1 03-1 (IMB/ K64), Muc100 05-7 (IMB/ K27; Przondovirus IL33), EKp2 09-52 (WRAIR/ K23; Przondovirus KpV763); OD600 reading of Appelmans cycle 1 for Kp strains 1 to 8 (Table 1); D) OD600 reading of Appelmans cycle 30 for Kp strains 1 to 8; E) Appelmans cocktail 30 phage isolated from Kp strain Li5.

Phages were subjected to propagation at an initial multiplicity of infection (moi) of 0.001 (moi 0.001: 10e4 Kp/well + 10 plaque forming unit (pfu) phage cocktail) on all eight hosts over multiple cycles in the Varioskan (LUX Multimode microplate reader; Thermo Fisher, Waltham, MA) at 37°C in 96-well plates overnight. After each cycle, products were pooled, centrifugated, and filtrated (0.2 µm).

Initial growth performance analyses indicated that the phages are individually lytic for their original hosts, but not to the five test hosts, non-susceptible Kp strains with additional restrictions (Figure 1C, Table 1). After 30 cycles, the final pool was tested on the eight target strains. The optical density (OD_600 nm_) curves indicated inhibitory growth effects compared to the controls in the original hosts as well as in all previously non-susceptible test hosts (Figure 1D and E). Cycle 30 cocktails from the eight strains were enriched on the respective strains and then further investigated by plaque assay.

Five phages were isolated from cell-free supernatant of cycle 30 cocktails (Table 1) and lytic phage genomes were sequenced. Four phages (Pin 21/0510-7, 9=11 and 13), represented the initial input phages (Figure 1B) without any nucleotide deviations in their genome sequences (data not shown). The fifth phage, however, lysed test host KpLi5, which was not lysed by any of the input phage. This turned out to be a novel induced temperate linear plasmid prophage *vB_tLPPP_KpLi5* (KpLi10-16) (Figure 1E). Its genome displays homology to N15/phiKO2-like telomere phages, which are plasmid prophages with covalently closed ends (hairpins), a prokaryotic system similar but distinct to eukaryotic poxviruses. Phages N15 (*E. coli*), phiKO2 (*K. oxytoca*), PY54 (*Yersinia enterocolitica*), VP882, VP58.5, VHML etc. (*Vibrio* species) are extrachromosomally resident in their hosts [9].

The 52,407 bp genome sequence of *vB_KpnS_KpLi5* (KpLi10-16) was deposited with Genbank under the accession number OQ830459. The phage genome could be confirmed as within the whole-genome sequence of the KpLi5 isolate, except for the chromosomal contig. The genomes represent all typical characteristics of the yet described enterobacterial telomere phages including the multifunctional replication protein (N15 RepA-like), the protelomerase (N15 TelN-like), and its telomere resolution site (N15 *telRL*-like). The sequence of the phages did not provide any hints for genes of undesired effects along therapeutical purposes, but undesired effects leading to changes of the growth performance of lysogenized bacteria, or their persistence, cannot be excluded [10, 11].

Phages derived from T7-like input phages with discrete genome mutations and expanded host range were not identified under the conditions used.

In summary, our controls showed re-isolation of input phages from our growth hosts. Interestingly, a novel temperate prophage was induced in our test host KpLi5 in the presence of all input phage, but not by any of them in separation. The underlying mechanisms of induction will be further analysed but may likely be a stress response to infection with multiple phages simultaneously. For the design of recombinant phage with extended host range for therapeutic purposes, we will redesign the experiment to offer a set of Kp hosts with a different set of capsule types.

Our results support linear use of phages in therapy, as recently advocated by the Turner lab [12], instead of simultaneous application in cocktails to limit the likelihood of stress inductions of possibly pathogenic temperate phages. Simultaneous use of phages in cocktails is still considered state of the art [13, 14]. Our findings may be useful to inform therapeutic approaches of our clinical partners [12-14], and the Phage4_1Health program [15]. (903)

## Statement of Author Contributions

Wrote first draft NJ, JJB

Wrote revisions: NJ, JH, SW, BS, AF, JJB

Scientific concept: JJB, BS, AF

Data evaluation: NJ, JH, JJ

## Statements and Declarations

The authors declare no competing interests.

## Acknowledgements and Funding

We thank Daniela Friese for technical assistance and Simone Eckstein for critical reading. This study was funded by the Medical Biological Defense Research Program of the Bundeswehr Medical Service. The work of JAH was supported by a grant (1322-820) of the BfR. Material has been reviewed by the Walter Reed Army Institute of Research. There is no objection to its presentation and/or publication. The opinions or assertions contained herein are the private views of the author, and are not to be construed as official, or as reflecting true views of the Department of the Army or the Department of Defense

